# Imd pathway-specific immune assays reveal NF-κB stimulation by viral RNA PAMPs in *Aedes aegypti* Aag2 cells

**DOI:** 10.1101/2020.06.30.179879

**Authors:** Tiffany A. Russell, Andalus Ayaz, Andrew D. Davidson, Ana Fernandez-Sesma, Kevin Maringer

## Abstract

**Background:** The mosquito *Aedes aegypti* is a major vector for the arthropod-borne viruses (arboviruses) chikungunya, dengue, yellow fever and Zika viruses. Vector immune responses pose a major barrier to arboviral transmission, and transgenic insects with altered immunity have been proposed as tools for reducing the global public health impact of arboviral diseases. However, a better understanding of virus-immune interactions is needed to progress the development of such transgenic insects. Although the NF-κB-regulated Toll and ‘immunodeficiency’ (Imd) pathways are increasingly thought to be antiviral, relevant pattern recognition receptors (PRRs) and pathogen-associated molecular patterns (PAMPs) remain poorly characterised in *A. aegypti*.

**Methodology/Principle Findings:** We developed novel RT-qPCR and luciferase reporter assays to measure induction of the Toll and Imd pathways in the commonly used *A. aegypti-derived* Aag2 cell line. We thus determined that the Toll pathway is not inducible by exogenous stimulation with bacterial, viral or fungal stimuli in Aag2 cells under our experimental conditions. We used our Imd pathway-specific assays to demonstrate that the viral dsRNA mimic poly(I:C) is sensed by the Imd pathway, likely through intracellular and extracellular PRRs. The Imd pathway was also induced during infection with the model insect-specific virus cricket paralysis virus (CrPV).

**Conclusions/Significance:** Our demonstration that a general PAMP shared by many arboviruses is sensed by the Imd pathway paves the way for future studies to determine how viral RNA is sensed by mosquito PRRs at a molecular level. Our data also suggest that studies measuring inducible immune pathway activation through antimicrobial peptide (AMP) expression in Aag2 cells should be interpreted cautiously given that the Toll pathway is not responsive under all experimental conditions. With no antiviral therapies and few effective vaccines available to treat arboviral diseases, our findings provide new insights relevant to the development of transgenic mosquitoes as a means of reducing arbovirus transmission.

**AUTHOR SUMMARY:** The mosquito *Aedes aegypti*, found globally across the tropics and subtropics, transmits viral diseases with a significant global public health impact, including chikungunya, dengue, yellow fever and Zika viruses. There are no antiviral drugs to treat these diseases and few effective vaccines. One way of reducing the global burden of mosquito-borne diseases would be to develop genetically modified mosquitoes unable to transmit viruses. One approach would be to alter the mosquitoes’ immune system to allow them to better fight viral infections. To do so, we first need to understand how viruses are detected by the mosquito immune system. We developed new methods of measuring immune responses in laboratory-cultured mosquito cells and used them to show that one specific arm of the immune system, called the ‘Imd pathway’, can detect the RNA that constitutes the genome of mosquito-borne viruses. These findings pave the way for future immune studies that could inform the development of transmission-incompetent mosquitoes. We also found that another arm of the immune system, called the ‘Toll pathway’, is not functional under any experimental conditions used in this study. This finding has implications for how different laboratories interpret data from these particular cultured cells.

## INTRODUCTION

Arthropod-borne viruses (arboviruses) are causing an ever-increasing global disease burden, with many arboviruses emerging, re-emerging or at risk of future emergence into the human population [1]. The mosquito species *Aedes aegypti* is distributed globally across the tropics and subtropics and is the most important arboviral vector of human diseases, including those caused by chikungunya, dengue, yellow fever and Zika viruses [2–4]. Given the lack of effective vaccines or antiviral therapies against most arboviruses, vector control measures that block transmission cycles remain one of the most effective ways of preventing human disease.

The ability of individual vector species and populations to transmit any given arbovirus is defined as ‘vector competence’ [5]. An improved understanding of the molecular virus-vector interactions underpinning vector competence would aid the development of genetically modified transmission-incompetent insects that could help reduce the global burden of arboviral disease. Mosquito immune responses have been shown to pose a major barrier to arbovirus transmission [6–12]. Therefore, manipulating immune responses has been proposed as an attractive approach for reducing the competence of *A. aegypti* for the broad range of arboviruses transmitted by this species [13].

Intracellular antiviral immune responses in mosquitoes include RNA interference (RNAi) pathways and inducible signalling cascades that result in the production of antimicrobial peptides (AMPs) and other gene products [14]. The expression of AMPs can be induced through the nuclear factor kappa-light-chain-enhancer of activated B cells (NF-κB)-regulated Toll and ‘immunodeficiency’ (Imd) pathways (Fig 1A), as well as the Janus kinase-signal transducer and activator of transcription (Jak-STAT) pathway [14]. In the model insect *Drosophila melanogaster*, which is classified in the order *Diptera* (flies) along with mosquitoes, AMPs like defensins have classically been considered to be Toll pathway-regulated while cecropins were considered to be Imd pathway-regulated [14]. However, the regulation of AMPs has been shown to be more complex than this [15,16], with species- and tissue-specific AMP regulation and cross-talk between NF-κB pathways [17–21].

**Fig 1.**
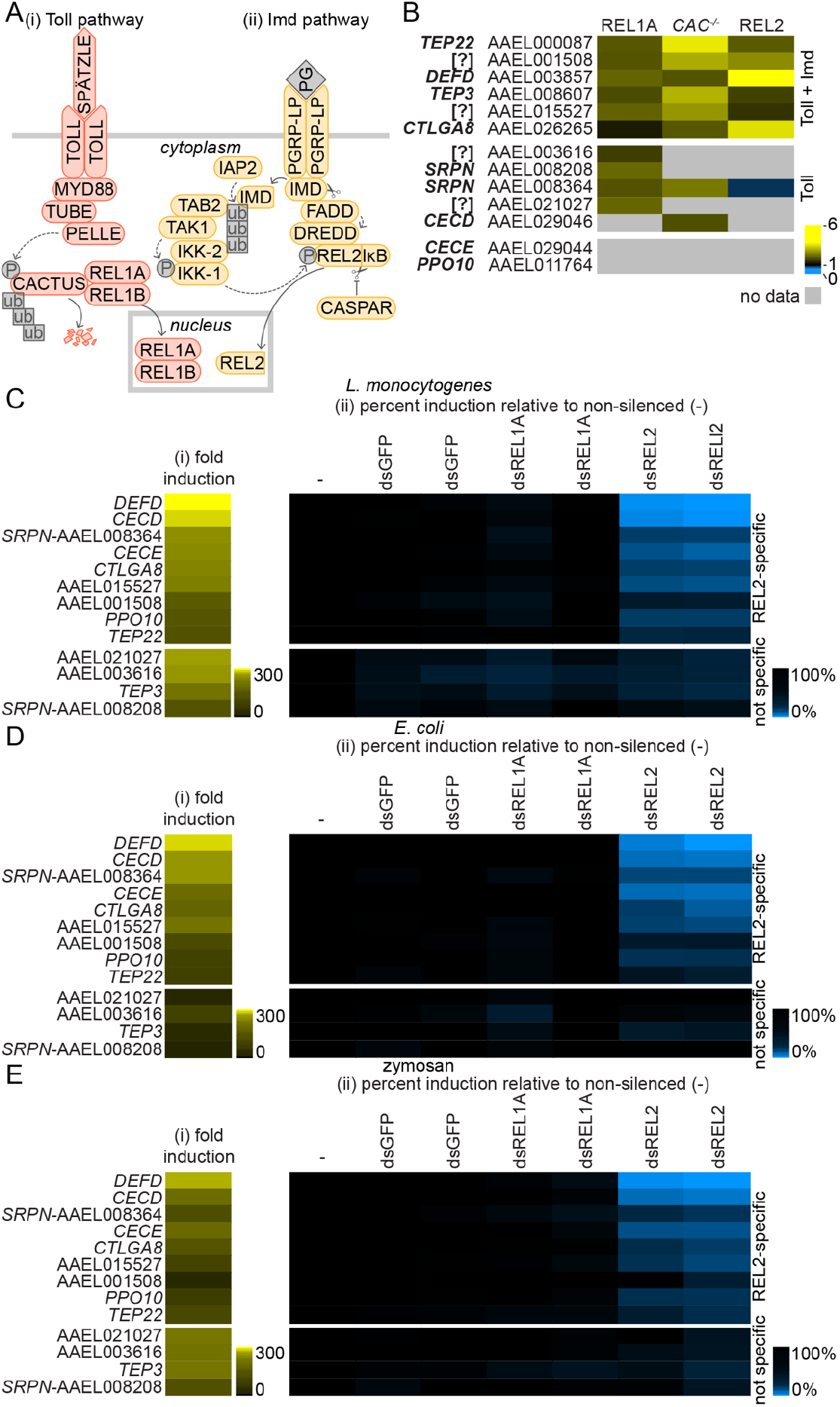
Stimulation of genes regulated through the Imd pathway by bacteria and zymosan in Aag2 cells. (A) Schematic of the *A. aegypti* Toll and Imd pathways. (i) The Toll pathway is induced when the transmembrane cytokine receptor TOLL binds to its dimeric cognate cytokine SPÄTZLE, which itself is activated by cleavage of the inactive precursor pro-SPÄTZLE following an extracellular proteolytic cascade initiated by PAMP detection. Subsequent TOLL dimerisation recruits the death domain-containing proteins myeloid differentiation primary response 88 (MYD88), TUBE and PELLE, resulting in phosphorylation and ubiquitin-mediated proteasomal degradation of the negative regulator CACTUS. This allows the nuclear translocation of the NF-κB-like transcription factor REL1A and its co-activator REL1B into the nucleus to induce the expression of AMPs and other immune-regulated genes. (ii) Imd pathway activation is associated with recruitment of the immunodeficiency (IMD) and Fas-associated death domain (FADD) proteins to activate the caspase DREDD (shown here following peptidoglycan (PG) sensing by the peptidoglycan recognition protein PGRP-LP). DREDD cleaves the transcription factor REL2 in a phosphorylation-dependent manner, allowing REL2’s N-terminal NF-κB domain to translocate into the nucleus and activate gene expression, while the C-terminal inhibitor of κB (IκB) domain remains cytoplasmic. REL2 is phosphorylated after caspases cleave IMD, allowing IMD to bind the inhibitor of apoptosis (IAP) family protein IAP2, an E3 ligase, which recruits and activates the IκB kinase (IKK) complex via K63 ubiquitin conjugation. CASPAR negatively regulates Imd pathway activation by inhibiting REL2 cleavage. (B) Summary of published gene expression data for transgenic *A. aegypti* overexpressing REL1A or REL2 [35], or wild-type *A. aegypti* intrathoracically inoculated with *CACTUS* dsRNA (CAC^-/-^) [7]. Gene expression data are replotted as fold induction relative to experimental control. Only genes relevant to the current study are shown. Genes are grouped based on induction through the Toll pathway, or both the Toll and Imd pathways. Gene annotation and Vectorbase Gene ID indicated; ‘[?]’ denotes hypothetical genes with no functional or protein identity annotation. (C-E) Fold induction (i) of genes measured by RT-qPCR in Aag2 cells stimulated for 24 h with 1,000 colony forming units (CFU)/cell of heat-inactivated *L. monocytogenes* (C) or *E. coli* (D), or 1 mg/ml zymosan (E). (ii) Percent gene induction in stimulated cells transfected with specified dsRNAs directed against *REL1A* (dsREL1A) or *REL2* (dsREL2) relative to the non-silenced control (‘-’), which was taken to be 100%. Two independent non-overlapping dsRNAs were tested for each gene; dsRNAs against *GFP* serve as a negative control. Log2 scale; genes consistently ordered by fold induction for *L. monocytogenes* (Ci) and REL2-specific or non-specific (‘not specific’) induction.

In *A. aegypti*, both the Toll and Imd pathways can be activated by arbovirus infection *in vivo* in a virus-specific manner [6–8,22–24]. Furthermore, NF-κB-regulated AMPs have been shown to possess antiviral activity [25,26]. Despite the growing evidence that NF-κB pathways play an antiviral role in mosquito vectors, a deep understanding of the viral pathogen-associated molecular patterns (PAMPs) and associated pattern recognition receptors (PRR) that stimulate NF-κB signalling is lacking. So far, NF-κB-regulated AMPs have been shown to be induced following detection of the dengue virus (DENV) envelope protein by complement-like thioester-containing proteins (TEPs) in combination with scavenger receptor C in *A. aegypti in vivo* [25]. In addition to this virus-specific PAMP-PRR combination, one may also expect more generally responsive PRRs to exist.

While vector competence studies can ultimately only be performed *in vivo*, cell culture models are invaluable tools for dissecting molecular virus-immune interactions in fine detail. Here, we developed highly specific RT-qPCR and luciferase reporter assays for measuring activation of the Imd pathway in the *A. aegypti*-derived Aag2 cell line. In the process, we demonstrated that Toll signalling is not activated by bacterial or viral PAMPs in Aag2 cells under our experimental conditions. Furthermore, the assays were used to show that the Imd pathway can be induced by viral infection and by poly(I:C), a mimic of viral dsRNA PAMPs. Our data suggest the existence of both cytoplasmic and cell membrane-localised PRRs capable of detecting viral RNA in *A. aegypti cells*.

## METHODS

### Cell cultures

Aag2 and S2 cells were a kind gift from Raul Andino (University of California, San Francisco, USA) and were maintained at 28°C in a humidified atmosphere without CO2 as previously described [27]. Culture medium was Leibovitz’s L-15 (Sigma-Aldrich, St. Louis, MO USA) supplemented with 10% (v/v) tryptose phosphate broth (Sigma-Aldrich), 10% (v/v) USA-origin foetal bovine serum (FBS), 2 mM L-glutamine, 0.1 mM non-essential amino acids, and 100 U/ml penicillin/100 μg/ml streptomycin (all ThermoFisher Scientific, Waltham, MA USA).

### Viruses

Cricket paralysis virus (CrPV) was a kind gift from Raul Andino. CrPV stocks were prepared by infecting confluent S2 cells one day post-seeding at a multiplicity of infection (MOI) of 0.2 in culture medium containing 2% (v/v) FBS and harvesting the culture supernatant four days post-infection. The supernatant was clarified by centrifugation and used as stock virus for infections. To determine the viral stock titre, S2 cells were seeded at 1 x 10^6^ cells/well in 12-well plates and infected one day post-seeding with serially diluted inoculum made up in PBS. After a one-hour incubation with gentle rocking at room temperature, an overlay of 0.8% (w/v) methyl cellulose (Sigma-Aldrich) in culture medium containing 2% FBS (v/v) was added. Cells were fixed after four days of culture at 28°C in 4% (w/v) paraformaldehyde and stained in 1% (v/v) crystal violet solution made up in 20% (v/v) ethanol.

### Bacteria and immune stimulation

*Escherichia coli* DH5α were purchased from ThermoFisher Scientific. *Listeria monocytogenes* and *Staphylococcus aureus* were kind gifts from Adolfo Garcia-Sastre and Flora Samaroo respectively (both Icahn School of Medicine at Mount Sinai, New York, NY USA). Bacteria were cultured overnight at 37°C with shaking in Luria Bertani medium (*E. coli*) or brain heart infusion broth (*L. monocytogenes, S. aureus*) without antibiotics, and were titrated on corresponding agar medium in 6-well plates. Culture media were from Sigma-Aldrich. Bacterial cultures were washed in phosphate buffered saline (PBS), resuspended in PBS and heat-inactivated at 60°C for 3 h (*E.coli, L. monocytogenes*) or 75°C for 6 h (*S. aureus*).

Cells were stimulated one day post-seeding by replacing the culture medium with fresh medium containing bacteria, zymosan or poly(I:C) at concentrations indicated. Zymosan A was purchased from Sigma-Aldrich and resuspended in sterile PBS. Poly(I:C) transfections were performed using Lipofectamine RNAi-MAX (ThermoFisher Scientific) one day post-seeding as per manufacturer’s instructions.

### Plasmids

pKM50, a constitutive *Renilla* luciferase expression plasmid, was previously described and is available through Addgene (addgene.org, Watertown, MA USA) with reference number 123656 [27]. A promoter-less vector backbone for generating mosquito firefly luciferase reporter constructs (pKM45; “luc” in Fig 3Bi) was cloned by replacing the Kozak sequence of pLUC-MCS (Agilent Technologies, Santa Clara, CA USA) with an *A. aegypti* Kozak sequence (GTGACC) and two additional TATA boxes following digestion with SacI and HindIII. pKM44 (“70-luc” in Fig 3Ai) was cloned similarly, with the addition of an upstream *D. melanogaster* minimal *HSP70* promoter element [28]. REL1A-specific (TCGAGACGC<u>GGGAAATTCC</u>AACG) and REL2-specific (CACGCTTTC<u>GGTGATTTAC</u>GCAC) NF-κB binding sites were characterised by Shin *et al*. [29]; the core NF-κB binding sites are underlined. Eight copies of each site were cloned into pKM44 or pKM45 following SacI and HindIII digestion to generate pKM51 (“1A-70-luc” in Fig 3Ai), pKM52 (“2-70-luc” in Fig 3Ai), pKM53 (“1A-luc” in Fig 3Bi) and pKM54 (“2-luc” in Fig 3Bi). pKM59 (“Mtk-wt-luc” in Fig 4Ai) was generated by inserting the 348 bp *Metchnikowin (Mtk*) enhancer element described by Senger *et al*. [30] into pKM45 following digestion with SacI and HindIII. REL1A-specific (pKM60, “Mtk-1A-1-luc” in Fig 4Ai) and REL2-specific (pKM62, “Mtk-2-luc” in Fig 4Ai) reporters were cloned using the same strategy, but the NF-κB binding sites were replaced with the same REL1A or REL2 binding sites used in the other reporters. The reporter containing five copies of the REL1A-specific *Mtk*-derived promoter (pAA3, “Mtk-1A-5-luc” in Fig 4Ai) was constructed sequentially by first introducing three copies of the modified enhancer element into pKM45 following digestion with SacI and HindIII to generate plasmid pAA1. We then sequentially generated pAA2 (four copies) and pAA3 (five copies) by inserting one addition copy of the modified enhancer element each time following digestion with XhoI and HindIII.

**Fig 2.**
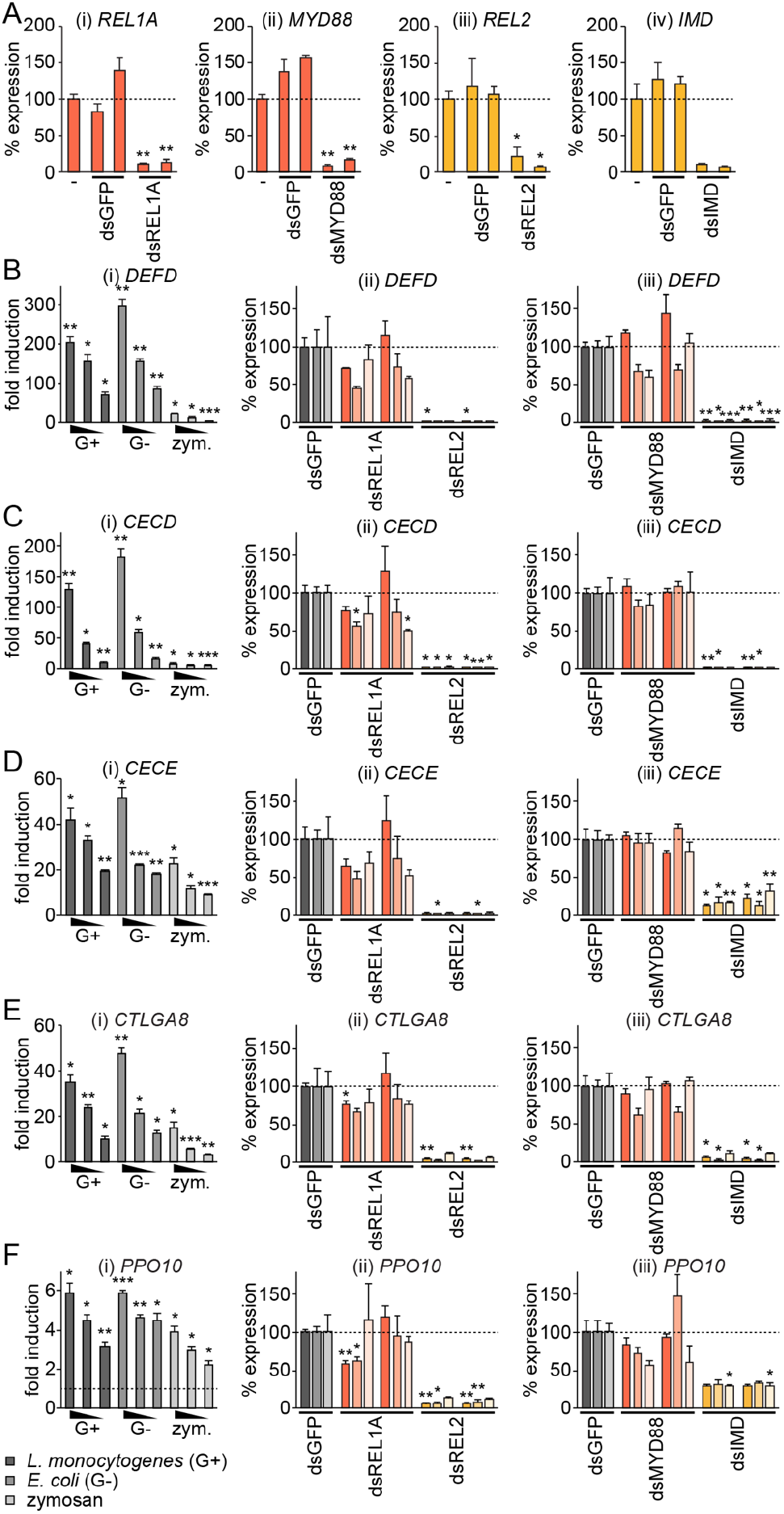
RT-qPCR assays specifically measuring Imd pathway activation in Aag2 cells. (A) Efficiency of gene silencing with two independent non-overlapping dsRNAs directed against *REL1A* (i), *MYD88* (ii), *REL2* (iii) or *IMD* (iv). Non-silenced cells transfected with no dsRNA (‘-’) taken as 100%; dsRNAs against GFP act as a negative control. (B-F) (i) Dose-dependent fold induction of indicated genes relative to unstimulated control in Aag2 cells 24 h post-stimulation with 1,000, 100 or 10 CFU/cell heat-inactivated *L. monocytogenes* (‘G+’) or *E. coli* (‘G-’), or 1, 0.1 or 0.01 mg/ml zymosan (‘zym.’). Dashed line in (F) indicates one-fold (no) induction. (ii, iii) Relative induction of indicated genes following stimulation with 1,000 CFU/cell heat-inactivated bacteria or 1 mg/ml zymosan in Aag2 cells previously transfected with indicated dsRNAs. Non-specific control dsRNA against *GFP* (dsGFP) taken as 100%; two independent non-overlapping dsRNAs were used for test gene knockdowns. Statistical significance for fold induction (i) or percent reduction (ii, iii) indicated; * *P* < 0.05; ** *P* < 0.01; *** *P* < 0.001. Error bars indicate standard error of the mean.

**Fig 3.**
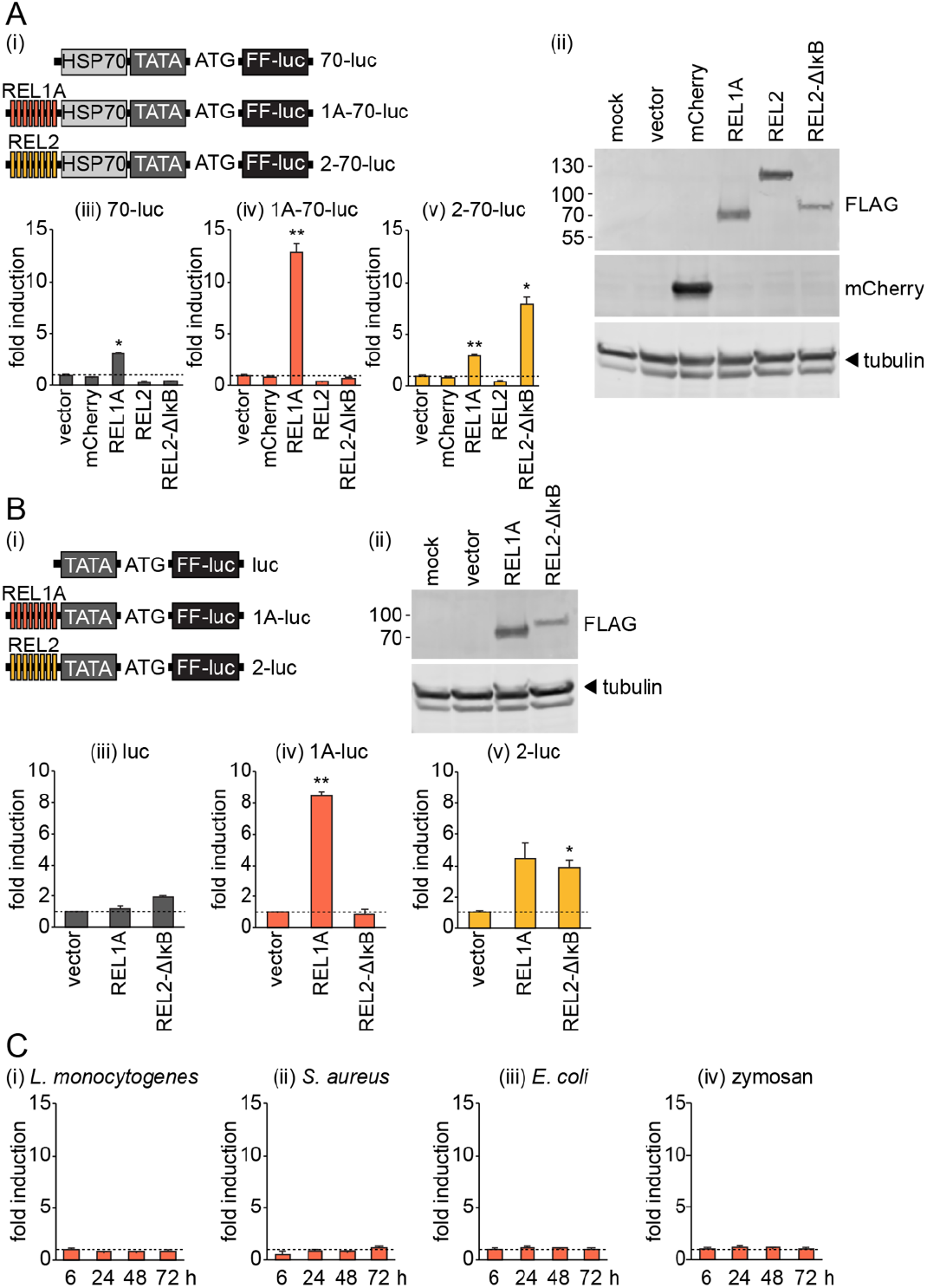
REL1A-specific NF-κB binding sites are non-responsive to exogenous immune stimulation in Aag2 cells. (A) (i) Schematic illustrating the promoter region of reporter plasmids expressing firefly luciferase (FF-luc) driven by eight copies of REL1A-specific (pKM51, ‘1A-70-luc’) or REL2-specific (pKM52, ‘2-70’luc’) NF-κB binding sites and the minimal *D. melanogaster HSP70* promoter element. TATA box and start codon (ATG) also shown. The control plasmid pKM44 (‘70-luc’) lacks NF-κB binding sites. (ii) Western blot confirming expression of mCherry and FLAG-tagged transcription factors in transiently transfected Aag2 cells. (iii-v) Fold induction of firefly luciferase from indicated reporter plasmids in Aag2 cells transiently transfected with mCherry or FLAG-tagged transcription factors relative to transfection with empty vector. (B) (i) Schematic illustrating plasmids generated as in (A) but lacking the minimal *HSP70* promoter element. luc, pKM45; 1A-luc, pKM53; 2-luc, pKM54. (ii) Western blot confirming expression of FLAG-tagged transcription factors in transiently transfected Aag2 cells. (iii-v) Firefly luciferase fold induction for indicated reporter plasmids in Aag2 cells transfected with FLAG-tagged transcription factors. (C) Firefly luciferase fold induction in Aag2 cells transfected with pKM51 (‘1A-70-luc’) and stimulated with 1,000 CFU/cell of heat-inactivated indicated bacteria or 1 mg/ml zymosan for the time periods stated. All firefly luciferase values were normalised to a co-transfected *Renilla* luciferase control plasmid. Statistical significance for fold induction indicated; * *P* < 0.05; ** *P* < 0.01. Error bars indicate standard error of the mean. Dashed line indicates one-fold (no) induction.

**Fig 4.**
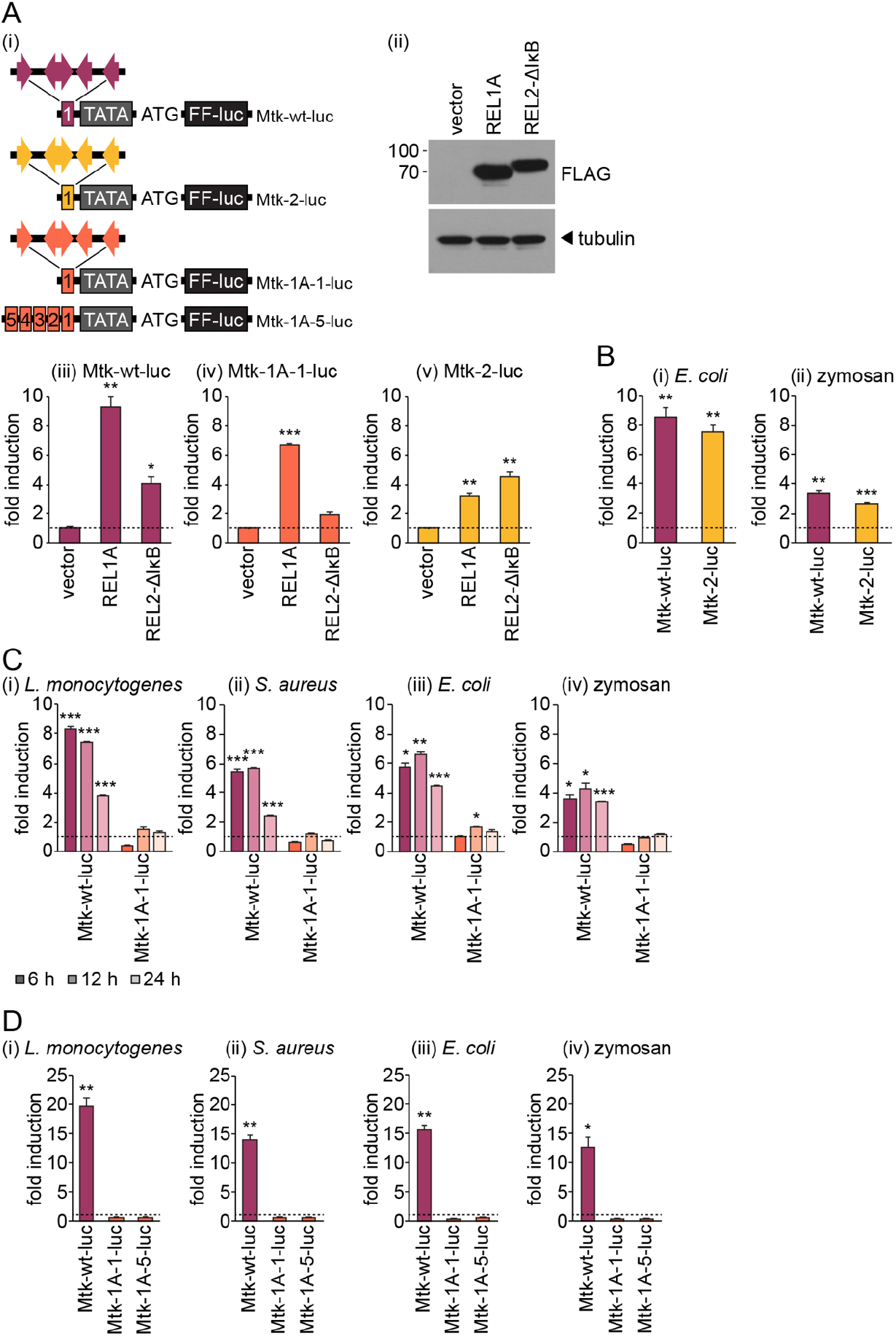
Substituting NF-κB binding sites for REL1A-specific sites renders the Toll and Imd pathway dually responsive *Mtk* promoter non-responsive to exogenous immune stimulation in Aag2 cells. (A) (i) Schematic illustrating the promoter region of reporter plasmids expressing firefly luciferase (FF-luc) driven by one copy of the wild-type *D. melanogaster Mtk* enhancer element (pKM59, ‘Mtk-wt-luc’), one copy of modified *Mtk* enhancer elements containing only REL2-(pKM62, ‘Mtk-2-luc’) or REL1A-specific (pKM60, ‘Mtk-1A-1-luc’) NF-κB binding sites, or five copies for the REL1A-specific *Mtk* enhancer element (pAA3, ‘Mtk-1A-5-luc’). TATA box and start codon (ATG) also shown, along with the orientation of NF-κB binding sites within the *Mtk* enhancer elements. (ii) Western blot confirming expression of FLAG-tagged transcription factors in transiently transfected Aag2 cells. (iii-v) Fold induction of firefly luciferase from indicated reporter plasmids in Aag2 cells transiently transfected with FLAG-tagged transcription factors relative to transfection with empty vector. (B) Firefly luciferase fold induction in Aag2 cells transfected with pKM59 (‘Mtk-wt-luc’) or pKM62 (‘Mtk-2-luc’) and stimulated with 1,000 CFU/cell of heat-inactivated *E. coli* (i) or 1 mg/ml zymosan (ii) for 24 h. (C) Firefly luciferase fold induction in Aag2 cells transfected with pKM59 (‘Mtk-wt-luc’) or pKM60 (‘Mtk-1A-1-luc’) and stimulated with 1,000 CFU/cell of indicated heat-inactivated bacteria or 5 mg/ml zymosan for the time periods stated. (D) Firefly luciferase fold induction in Aag2 cells transfected with pKM59 (‘Mtk-wt-luc’), pKM60 (‘Mtk-1A-1-luc’) or pAA3 (‘Mtk-1A-5-luc’) and stimulated with 1,000 CFU/cell of indicated heat-inactivated bacteria or 1 mg/ml zymosan for 24 h. All firefly luciferase values were normalised to co-transfected *Renilla* luciferase control plasmid. Statistical significance for fold induction indicated; * *P* < 0.05; ** *P* < 0.01; *** *P* < 0.001. Error bars indicate standard error of the mean. Dashed line indicates one-fold (no) induction.

The mCherry expression vector pKM7 was constructed by digesting pIEx-EGFP [31], a kind gift from Doug Brackney (The Connecticut Agricultural Experiment Station, New Haven, CT USA), with XhoI and NcoI and substituting the *EGFP* sequence for the *mCherry* sequence. pKM28 encodes *A. aegypti* REL1A with a C-terminal FLAG tag (GATTACAAGGATGACGATGACAAG, peptide sequence DYKDDDD), and was cloned by digesting pIEx-EGFP with XhoI and NcoI and replacing the *EGFP* sequence with the FLAG-tagged *REL1A* sequence (vectorbase.org reference AAEL007696). pKM5 encodes *A. aegypti* REL2 with an N-terminal FLAG tag, which was cloned in the same way as pKM28 using the *REL2* (AAEL007624) sequence. There are three isoforms of REL2 in *A. aegypti* produced by alternative splicing [32]; we cloned the dominant full-length sequence (NCBI reference XM_001658467.1), not the ‘Rel-type’ lacking the C-terminal IκB domain or the ‘IκB-type’ lacking much of the N-terminus. To generate the plasmid expressing constitutively active N-terminally FLAG-tagged REL2 lacking its IκB domain (pKM33), we amplified the N-terminal sequence of R EL2 up to the caspase cleavage site from pKM5 and cloned it into pIEx-EGFP as for the other expression vectors. The caspase cleavage site was determined to be 552LETD555 by reference to *D. melanogaster* RELISH and by using http://www.casbase.org/casvm/index.html [33]. All *A. aegypti* gene sequences were PCR amplified from Aag2 cells. We also generated a control vector expressing just the FLAG tag (pKM3) by substituting the *EGFP* from pIEx-EGFP with the FLAG sequence following digestion with XhoI and NcoI.

All cloning was performed using In-Fusion cloning (Takara Biosciences, Mountain View, CA USA). Insert sequences were purchased from Sigma-Aldrich or Integrated DNA Technologies (Coralville, IA USA). All of the plasmids that we generated here have been deposited with Addgene with reference numbers 154152-154168.

### RNAi gene silencing

Aag2 cells were seeded at 1 × 10^5^ cells/well in 96-well plates and concurrently transfected at a final dsRNA concentration of 10 nM using Lipofectamine RNAi-MAX (ThermoFisher Scientific) as per manufacturer’s instructions. dsRNA was prepared by PCR amplification with primers detailed in Table 1 using RNA purified from Aag2 cells as a template; *EGFP* dsRNA was prepared using plasmid pIEx-EGFP as a template. PCR amplicons were subjected to *in vitro* transcription using the MEGAshortscript *in vitro* transcription kit (ThermoFisher Scientific) and gel purified before storage in aliquots at −80°C.

**Table 1.**
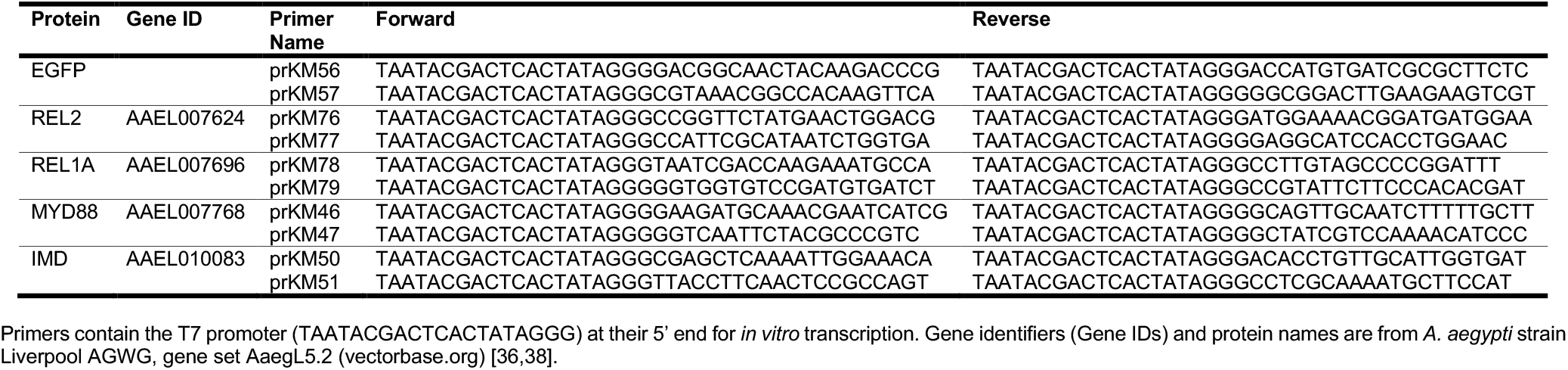
dsRNA Primers.

### RT-qPCR

Total RNA was extracted from 1 × 10^5^ cells using the Quick-RNA MiniPrep kit with Zymo-Spin IIICG columns (Zymo Research, Irvine, CA USA) as per manufacturer’s instructions. cDNA was generated from RNA using the iScript reverse transcription supermix for RT-qPCR (Bio-Rad, Hercules, CA USA) as per the manufacturer’s instructions. qPCR amplification and detection was performed using the Hot FIREPol EvaGreen qPCR supermix Plus, no ROX (Solis Biodyne, Tartu, Estonia) and primers detailed in Table 2 at 95°C for 10 min, followed by 40 cycles of 95°C for 15 s and 60°C for 30 s, as per manufacturer’s instructions on a CFX96 Touch Real-Time PCR Detection System (Bio-Rad). Gene expression was normalised against the 40S ribosomal protein S7 (RPS7) mRNA.

**Table 2.**
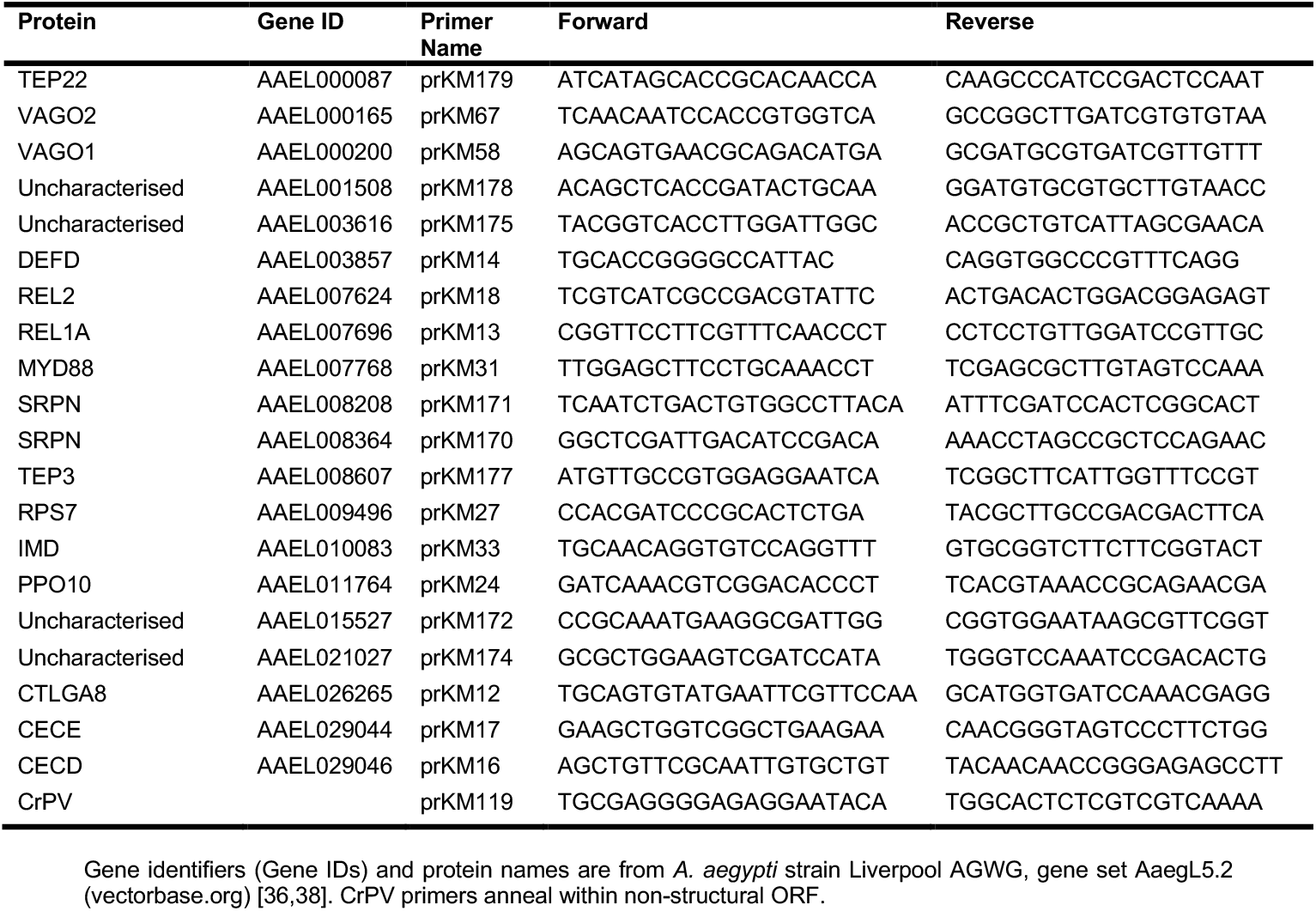
RT-qPCR Primers.

CrPV RNA standards for absolute quantification of viral RNA were generated by PCR amplifying nucleotides 2,173-3,200 of the CrPV genomic RNA non-structural ORF from infected Aag2 cells using the following primers: prKM120F, TAATACGACTCACTATAGGGGTTCTCCTCTCTCCTGTCAAAG; prKM120R, TCACTAATAAATTTGTAAGCTTTGTTCTTAA. The forward primer contains the T7 promoter sequence at its 5’ end to generate a positive-sense RNA standard. PCR was performed using the SuperScript III One-Step RT-PCR System with Platinum Taq DNA Polymerase (ThermoFisher Scientific) as per manufacturer’s instructions with the addition of 2 μl MgSO4, 2.5 μl bovine serum albumin (BSA) (New England Biolabs, Ipswich, MA USA) and 2.5 μl dimethyl sulfoxide (DMSO) (Sigma-Aldrich) in a 25 μl reaction volume. Cycles were 55°C for 30 min, 94°C for 2 min, followed by 40 cycles of 94°C for 30 s, 59°C for 30 s, 68°C for 4 min, and a final extension at 68°C for 10 min. Gel purified cDNA was subjected to *in vitro* transcription using the MEGAscript T7 High Yield Transcription Kit (ThermoFisher Scientific) as per manufacturer’s instructions. RNA standards were treated with the DNA-free DNA removal kit (ThermoFisher Scientific) as per manufacturer’s instructions and gel purified before use.

### Western blots

Aag2 cells were seeded at 1 × 10^6^ cells/well in 12-well plates and transfected at the time of seeding with 3 μg protein expression plasmid using Lipofectamine 3000 transfection reagent (ThermoFisher Scientific) as per manufacturer’s instructions. Cells were harvested 48 h post-seeding/-transfection. Cells were washed once with PBS and resuspended in 2X SDS-PAGE buffer consisting of 10 mM Tris, 4% (w/v) sodium dodecyl sulphate (SDS), 10% (v/v) β-mercaptoethanol, 20% (v/v) glycerol and 0.2% (w/v) bromophenol blue (all reagents from Sigma Aldrich). Samples were boiled for 3 min and sonicated for 30 s prior to loading on 4-20% Mini-PROTEAN TGX precast protein gels (Bio-Rad) and electrophoresis in Tris-glycine buffer. Gels were transferred to polyvinylidene fluoride membranes, blocked overnight in TBS-Tween with 5% (w/v) milk, and incubated with the following primary antibodies as indicated: mouse anti-mCherry ‘1C51’ (Abcam, Cambridge, UK; catalogue number ab125096), mouse anti-FLAG ‘M2’ (Sigma-Aldrich; F3165), rabbit anti-tubulin (alpha) ‘EP1332Y’ (Abcam; ab18251). Membranes were then incubated with IRDye 800CW goat anti-mouse IgG (H+L) or IRDye 680RD goat anti-rabbit IgG (H+L) secondary antibodies (LI-COR Biosciences, Lincoln, NE USA) as appropriate. Blots were visualised and analysed using the Odyssey CLx imaging system (LI-COR Biosciences).

### Luciferase assays

Aag2 cells were seeded at 1 × 10^5^ cells/well in 96-well plates and transfected at the time of seeding with 10 ng pKM50 (*Renilla* luciferase transfection control) and 50 ng/well of the respective firefly reporter plasmid using Lipofectamine 3000 transfection reagent (ThermoFisher Scientific) as per manufacturer’s instructions. Where appropriate, 300 ng/well protein expression plasmid was also included. Cells were harvested in passive lysis buffer (Promega, Madison, WI USA) and incubated with gentle agitation for 20 min at room temperature. Firefly and *Renilla* luciferase activity were measured using a CLARIOstar plate reader (BMG Labtech, Aylesbury, UK) and home-made luciferase assay reagents as published [34].

### Immune gene data mine and transcription factor binding site analysis

Genes overexpressed compared to wild-type mosquitoes in transgenic *A. aegypti* overexpressing REL1A or REL2 were taken from Zou *et al*. supplemental tables S1, S2, S3 and S4 [35]. Genes upregulated in *A. aegypti* intrathoracically injected with *CACTUS* dsRNA were taken from Xi *et al*. supplemental table S6 [7]. Data were pooled and at the time of the analysis gene identifiers were converted to *A. aegypti* strain Liverpool AGWG genome assembly AaegL3 (vectorbase.org) [36,37]. Throughout this manuscript we use the most up-to-date gene identifiers from gene set AaegL5.2 [36,38].

NF-κB binding sites are enriched in the 500 bp upstream of the 5’UTR in *D. melanogaster* [39]. Therefore, for genes with an annotated 5’UTR in the AaegL3 assembly, the 1,000 bp upstream and the first 100 bp downstream of the transcription start site were downloaded from vectorbase.org to provide maximum opportunity for identifying NF-κB binding sites. For AAEL003857 (*DEFD*), AAEL011764 (*PPO10*), AAEL026265 (*CTLGA8*) and AAEL029044 (*CECE*) the promoters were not annotated at the time of the analysis. For these genes, the 2,000 bp upstream and the 100 bp downstream of the start codon were downloaded from vectorbase.org, as our own analysis of 129 genes with annotated 5’UTRs found that this would capture the 500 bp upstream of the 5’UTR in 98% of cases. Sequences in FASTA format were analysed for the presence of predicted *D. melanogaster* NF-κB binding sites using MatInspector (genomatix.de, Genomatix, Munich, Germany). In addition, we searched for *A. aegypti* REL1A- and REL2-specific and dually responsive NF-κB binding sites defined by Shin *et al*. [29].

### Data analysis and images

All data were analysed in Microsoft Excel (Microsoft Corporation, Redmund, WA USA). Statistical significance was tested using homoscedastic Student’s *t* test assuming unequal variance. The Bonferroni correction was applied for multiple comparisons. All images were prepared in Adobe Illustrator (Adobe Systems, San Jose, CA USA). Images were cropped and annotated for clarity only.

## RESULTS

### Immune-regulated genes are activated via the Imd but not the Toll pathway in Aag2 cells

To facilitate studies into molecular virus-immune interactions, we set out to develop RT-qPCR assays to measure the activation of the Imd and Toll pathways in Aag2 cells. We selected genes previously shown to be upregulated *in vivo* in *A. aegypti* by endogenous activation of NF-κB signalling, either through the transgenic overexpression of the transcription factors REL1A (Toll pathway) or REL2 (Imd pathway), or by transient dsRNA silencing of the negative regulator CACTUS (Toll pathway) (Fig 1B) [7,35]. Published data on Imd pathway activation via transient dsRNA silencing of the negative regulator CASPAR were not included in the analysis, as the authors of the original studies acknowledged that there was little overlap with REL2 overexpression data [7,35]. While components within immune pathways can be used as markers of pathway activation, these tend to be induced to a lesser degree than downstream secreted effectors [40] and were therefore excluded. In total we selected fourteen genes from these data sets for validation in Aag2 cells, including one defensin (*DEFD*), two cecropins (*CECD*, *CECE*), two serpins (currently annotated simply as *‘SRPN’*), two TEPs (*TEP3, TEP22*), one prophenoloxidase (*PPO10*), one C-type lectin (galactose-binding) (*CTLGA8*), and four ‘hypothetical’ genes with no functional or identity annotation (AAEL001508, AAEL003616, AAEL015527, AAEL021027) (Fig 1B). This panel of genes was chosen from our data mine to represent a broad range of proteins with known immune functions; CECE and PPO10 were selected for their known immune functions despite not being represented in the published *in vivo* data. Throughout this manuscript we refer to all genes by their current annotation and gene identifiers on vectorbase.org (*A. aegypti* strain Liverpool AGWG, gene set AaegL5.2) [36,38]. We also checked for the presence of predicted NF-κB binding sites in the promoters of these genes (Table 3).

**Table 3.** Predicted NF-κB binding sites in the promoters of immune-regulated genes.

| Protein | Gene ID | AaREL1A | AaREL2 | AaREL1A + AaREL2 | DmNF-κB |
| --- | --- | --- | --- | --- | --- |
| TEP22 | AAEL000087 | -334 (+)<br>-511 (+)<br>-689 (-) |  | -952 (-) | -226 (+)<br>-686 (+) |
| Uncharacterised | AAEL001508 | -123 (-)<br>-940 (-) |  | +6 (-)<br>-68 (-)<br>-686 (+) | -120 (+)<br>-540 (+)<br>-937 (+)<br>-967 (+) |
| Uncharacterised | AAEL003616 |  |  |  | -281 (+)<br>-307 (+)<br>-397 (+)<br>-471 (+) |
| SRPN | AAEL008208 | -17 (-)<br>-26 (-)<br>-35 (-)<br>-45 (-)<br>-63 (-) |  |  | -14 (+)<br>-23 (+)<br>-32 (+)<br>-60 (+)<br>-61 (+)<br>-90 (+)<br>-299 (-)<br>-825 (-) |
| SRPN | AAEL008364 | -113 (+) |  |  | -11 (-)<br>-836 (+)<br>-980 (+) |
| TEP3 | AAEL008607 | -22 (+)<br>-31 (-)<br>-138 (-) | -48 (+)<br>-49 (-)<br>-173 (+) | -145 (-)<br>-173 (-)<br>-263 (-) | -33 (+)<br>-34 (+)<br>-171 (-) |
| SRPN | AAEL015527 | +42 (-)<br>-238 (-)<br>-523 (+)<br>-715 (-)<br>-803 (-) |  |  | -235 (+)<br>-493 (-)<br>-520 (-) |
| Uncharacterised | AAEL021027 | +17 (+) |  |  | +20 (-)<br>-98 (-)<br>-544 (+)<br>-947 (-) |
| CECD | AAEL029046 | -959 (-) |  | -589 (+)<br>-803 (-)<br>-837 (-) | -199 (+)<br>-561 (-)<br>-576 (-)<br>-864 (+) |

Gene identifiers (Gene IDs) are from *A. aegypti* strain Liverpool AGWG, gene set AaegL5.2 (vectorbase.org) [36,38]. AaREL1A, sites responsive to *A. aegypti* REL1A; AaREL2, sites responsive to *A. aegypti* REL2; AaREL1A + AaREL2, sites responsive to both REL1A and REL2 of *A. aegypti* [29]. DmNF-κB, sites responsive to *D. melanogaster* NF-κB. '-' denotes site within 1,000 bp upstream of 5'UTR; '+' denotes site within first 100 bp of 5'UTR; '(+)' denotes site on sense strand; '(-)' denotes site on antisense strand. For AAEL003857 (*DEFD*), AAEL011764 (*PPO10*), AAEL026265 (*CTLGA8*) and AAEL029044 (*CECE*) the promoters were not annotated at the time of the analysis, and no NF-κB sites were identified within 2,000 bp upstream of the ATG.

Next, we tested the induction of the selected genes by RT-qPCR following treatment with heat-inactivated Gram-positive (*Listeria monocytogenes*) or Gram-negative (*Escherichia coli*) bacteria or the fungal PAMP zymosan, which are classic stimuli of insect NF-κB pathways [14]. The genes were induced to a greater or lesser extent by each of the stimuli, with up to 300-fold induction seen for some genes (Fig 1Ci, Di, Ei). Although several other studies have measured the expression of various immune-regulated genes by RT-qPCR in Aag2 cells [40–42], only one previous study validated which specific immune pathways these genes were regulated by, with validation only performed for stimulation with heat-inactivated *E. coli* [42]. We therefore tested the impact that transient silencing of *REL1A* or *REL2* with dsRNA had on the induction of our putative NF-κB-regulated genes following stimulation with our panel of heat-inactivated bacteria or zymosan. Of the genes tested, nine were activated to markedly lower levels with all stimuli following silencing of *REL2*, while no impact was observed following *REL1A* silencing (*DEFD, CECD, CECE*, AAEL001508, AAEL008364 (*SRPN*), AAEL015527, *CTLGA8, PPO10, TEP22*) (Fig 1Cii, Dii, Eii; knockdown efficiency shown in Fig 2Ai, 2Aiii). These genes thus appear to be regulated by the Imd pathway in Aag2 cells. Four genes (AAEL003616, AAEL008208 (*SRPN*), AAEL021027, *TEP3*) were not clearly regulated by either pathway.

To validate a subset of the apparently Imd pathway-regulated genes in more detail, we verified the dose-dependency of the induction of *DEFD, CECD, CECE, CTLGA8* and *PPO10* (Fig 2Bi, Ci, Di, Ei, Fi). Both Gram-positive and Gram-negative bacteria stimulated robust expression of these genes, with up to 200- to 300-fold induction for *DEFD* and *CECD*, and approximately 40- to 50-fold induction for *CECE* and *CTLGA8*. We consistently saw lower induction for *PPO10*, and lower induction of all genes with zymosan compared to heat-inactivated bacteria. To more extensively test the NF-κB pathways through which these genes were induced, we repeated the stimulation in the presence of dsRNA targeting *REL1A, REL2, MYD88* or *IMD*, with MYD88 and IMD acting earlier in the Toll and Imd signalling cascades respectively compared to the transcription factors REL1A and REL2. Once again, we consistently found that all genes were induced solely through the Imd pathway for all stimuli tested (Fig 2). Our findings are in line with recently published data by Zhang *et al*. that also found little effect of transient *REL1A* or *MYD88* silencing on the induction of immune genes by *E. coli* in Aag2 cells [42].

### REL1A-specific transcription factor binding sites are non-responsive in Aag2 cells

Given that we and others [42] only tested a limited number of genes and that there could still feasibly exist other Toll pathway-inducible genes in Aag2 cells, we turned to luciferase reporters to further investigate the responsiveness of REL1A-specific transcription factor binding sites in Aag2 cells. NF-κB reporters previously used in Aag2 cells [23] were based on *D. melanogaster* promoters that may not faithfully recapitulate the regulation of *A. aegypti* promoters. We therefore set out to develop NF-κB reporters based on transcription factor binding sites previously validated in *A. aegypti* [29].

First, we constructed plasmids in which firefly luciferase expression is driven by eight copies of REL1A- or REL2-specific NF-κB binding sites upstream of the *D. melanogaster* minimal *HSP70* promoter element [28], which is present in the promoter of many insect expression plasmids (Fig 3Ai). We co-transfected Aag2 cells with each reporter and plasmids encoding mCherry or FLAG-tagged *A. aegypti* REL1A, REL2 or a truncated constitutively active REL2 construct lacking the inhibitory IκB domain (Fig 3Aii). None of the reporter plasmids were activated by the mCherry negative control (Fig 3Aiii-v). While the REL1A-specific reporter (1A-70-luc) was robustly induced specifically by REL1A, the REL2-responsive reporter (2-70-luc) was induced both by the REL2-ΔIκB construct and by REL1A (Fig 3Aiv, v). Since the control reporter lacking NF-κB binding sites (70-luc) was also induced by REL1A overexpression (Fig 3Aiii), we initially presumed this to be due to the activation of the minimal *DmHSP70* promoter by REL1A. Only the REL2-△IκB construct and not full-length REL2 activated the REL2-responsive reporter (2-70-luc) (Fig 3Av), presumably because the presence of the IκB domain prevented the nuclear translocation of full-length REL2 in the absence of exogenous immune stimulation.

To overcome the presumed non-specific activation of the REL2-responsive reporter (2-70-luc) by REL1A, we generated equivalent reporters lacking the minimal *DmHSP70* promoter (Fig 3Bi). Although the control reporter lacking NF-κB binding sites (luc) no longer responded to REL1A overexpression, and the REL1A-specific reporter (1A-luc) remained specific to REL1A stimulation, the REL2-responsive reporter (2-luc) was still induced by both REL1A and REL2-ΔIκB overexpression (Fig 3B). We therefore conclude that the previously reported REL2-specific NF-κB binding sites are not REL2-specific in Aag2 cells. We note that our reporters lacking the minimal *DmHSP70* promoter (Fig 3B) were less inducible than the original reporters (Fig 3A). We therefore used our original REL1A-specific reporter containing the minimal *DmHSP70* promoter (1A-70-luc) to test its inducibility by heat-inactivated Gram-positive (*L. monocytogenes*, *Staphylococcus aureus*) or Gram-negative (*E. coli*) bacteria or zymosan. In line with our RT-qPCR data (Fig 1, 2), we observed no Toll pathway activation by any of the tested stimuli using this luciferase reporter (Fig 3C). Indeed, we tested reporter induction over a range of times post-stimulation to exclude the possibility that our previous experiments had missed a potentially transient Toll pathway activation.

Finally, we sought to exclude the possibility that our REL1A-specific reporters (1A-70-luc, 1A-luc) were non-responsive due to the lack of binding sites for other transcription factors required for co-stimulation of the Toll pathway. The *D. melanogaster Metchnikowin (Mtk*) promoter is regulated independently by both the Toll and Imd pathways [43,44] and contains four NF-κB binding sites along with binding sites for other transcription factors that co-stimulate NF-κB signalling [30]. We generated a firefly luciferase reporter driven by one copy of a previously defined 348 bp enhancer region from the *D. melanogaster Mtk* promoter (Fig 4Ai) [30]. We also generated additional reporters in which the NF-κB binding sites in the *Mtk* enhancer were replaced with the same REL1A- or REL2-specific NF-κB binding sites used in our other reporters (Fig 4Ai). We first confirmed that the wild-type *Mtk* reporter (Mtk-wt-luc) was activated by both REL1A and REL2-ΔIκB overexpression (Fig 4Aii, iii). Replacing the NF-κB binding sites with REL1A-specific NF-κB binding sites (Mtk-1A-1-luc) made the reporter REL1A-specific (Fig 4Aiv). In agreement with our previous data (Fig 3), the *Mtk* reporter containing only ‘REL2-specific’ NF-κB binding sites (Mtk-2-luc) was activated by both REL1A and the REL2-ΔIκB construct (Fig 4Av). Nevertheless, we used this reporter to confirm that our manipulation of the *Mtk* enhancer region did not have a notable impact on the activation of Imd pathway-responsive reporters by *E. coli* or zymosan treatment (Fig 4B). In contrast, substituting the NF-κB binding sites for exclusively REL1A-specific NF-κB binding sites (Mtk-1A-1-luc) caused the reporter to become non-responsive to any of the immune stimuli tested (Fig 4C). We furthermore determined that even a reporter containing five copies of the REL1A-specific *Mtk* enhancer (Mtk-1A-5-luc) still failed to respond to immune stimulation (Fig 4Ai, D).

In summary, we conclude that under our experimental conditions the Toll pathway in Aag2 cells is not responsive to stimulation with a number of well-established immune stimuli. Therefore, our system is exquisitely specific for the Imd pathway, allowing the interactions of viruses with the Imd pathway to be investigated with high confidence.

### Activation of REL2-regulated genes by viral PAMPs in Aag2 cells

We next investigated whether the expression of REL2-regulated genes in *A. aegypti* cells is activated by stimulation with dsRNA, a PAMP commonly found in cells infected with RNA viruses [45]. Of the five genes that we validated as REL2-inducible (*DEFD*, *CECD*, *CECE, CTLGA8, PPO10*), all were induced in a dose-dependent manner 24 h post-transfection with poly(I:C) (Fig 5A). Given that dsRNA is likely to be detected through different PRRs than bacterial or fungal PAMPs, we verified that poly(I:C) signals through REL2 using our wild-type and REL1A-responsive *Mtk* promoter reporters (Fig 5B). Similarly to bacteria and zymosan (Fig 4C and D), poly(I:C) induced luciferase expression from the wild-type, but not the REL1A-responsive, reporter, indicating that poly(I:C) is not activating the Toll pathway under our experimental conditions. Interestingly, poly(I:C) stimulated the expression of REL2-regulated genes both when it was transiently transfected into Aag2 cells and when it was simply added to the culture medium (Fig 5C). However, the levels of gene induction were up to twenty-fold greater during transient transfection. While we found that there was some variability in the levels of REL2-regulated gene induction between genes and experiments, the observation that poly(I:C) induces REL2-regulated genes itself was highly reproducible.

**Fig 5.**
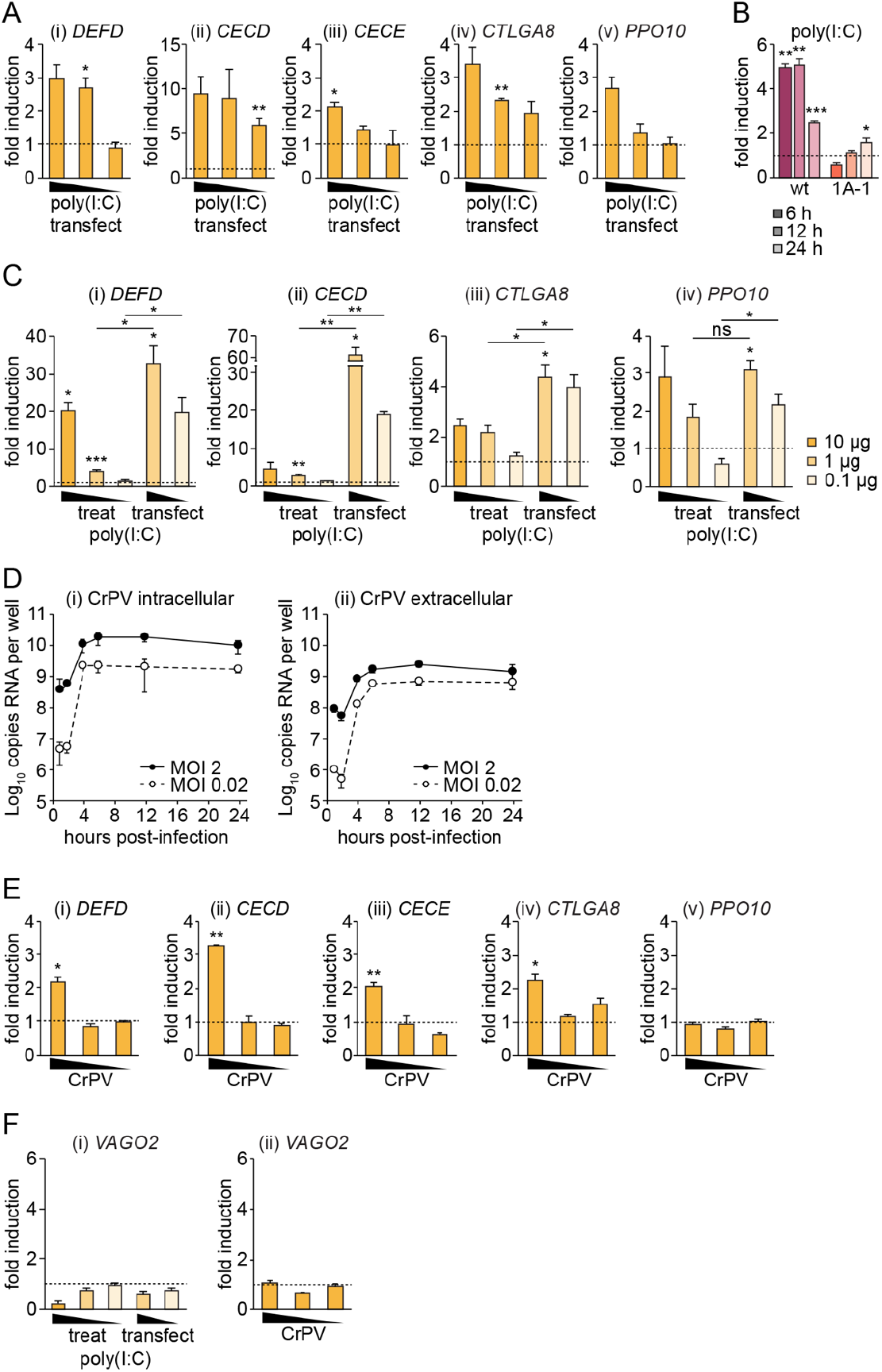
Induction of the Imd pathway in Aag2 cells stimulated with poly(I:C) or infected with CrPV. (A) Fold induction of indicated genes measured by RT-qPCR in Aag2 cells transiently transfected for 24 h with 10, 1 or 0.1 μg/well poly(I:C) in 24-well plates. (B) Firefly luciferase fold induction in Aag2 cells grown in 96-well plates and transfected with *Mtk*-based wild-type (pKM59, ‘wt’) or REL1A-specific (pKM60, ‘1A-1’) firefly reporter plasmids prior to transfection with 10 μg/well poly(I:C) for the time periods stated. Firefly luciferase values were normalised to a co-transfected *Renilla* luciferase control plasmid. (C) Fold induction of indicated genes measured by RT-qPCR in Aag2 cells transiently transfected with 1 or 0.1 μg/well poly(I:C) or treated with 10, 1 or 0.1 μg/well poly(I:C) for 24 h in 24-well plates. (D) Viral growth curves showing absolute quantification of total intracellular (i) or extracellular (ii) viral RNA copies per well in Aag2 cells infected with CrPV in 24-well plates. (E) Fold induction of indicated genes measured by RT-qPCR in Aag2 cells infected with CrPV at MOI 2, 0.2 or 0.02 for 24 h. (F) Fold induction of *VAGO2* measured by RT-qPCR in Aag2 cells transiently transfected with 1 or 0.1 μg/well poly(I:C) or treated with 10, 1 or 0.1 μg/well poly(I:C) for 24 h in 24-well plates (i), or infected with CrPV at MOI 2, 0.2 or 0.02 for 24 h (ii). Statistical significance for fold induction, or relative differences in induction levels (C, bars), indicated; * *P* < 0.05; ** *P* < 0.01; *** *P* < 0.001; ns, non-significant. Error bars indicate standard error of the mean, except in (D) where error bars indicate standard deviation. Dashed line indicates one-fold (no) induction.

To establish whether REL2-regulated genes can also be induced during RNA virus infection, we tested a number of viruses for their ability to infect Aag2 cells and induce AMP expression. We found that the insect-specific dicistrovirus virus cricket paralysis virus (CrPV), which is often used as a model virus in immune studies in *D. melanogaster*, replicates with rapid growth kinetics in Aag2 cells (Fig 5D). We detected the accumulation of CrPV RNA both intracellularly and extracellularly at high and low MOI, peaking by 6 hours post-infection, suggesting that CrPV replicates in Aag2 cells and releases viral particles into the extracellular medium. Furthermore, we detected the induction of *DEFD, CECD, CECE* and *CTLGA8*, but not *PPO10*, during CrPV infection (Fig 5E).

VAGO has been reported to be a virus-inducible cytokine regulated through REL2 that is involved in activating innate immune signalling in *D. melanogaster* and mosquitoes [46–49]. There are two *VAGO* genes in *A. aegypti, VAGO1* (AAEL000200) and *VAGO2* (AAEL000165) [49]. Despite extensive experiments and the reproducible induction of other REL2-regulated immune genes, we failed to detect the induction of *VAGO2* during CrPV infection or stimulation with poly(I:C) (Fig 5F). We were unable to detect the expression of *VAGO1* in either stimulated or unstimulated Aag2 cells.

## DISCUSSION

We developed a comprehensive and extensively characterised set of RT-qPCR and firefly luciferase assays to measure signalling through the Imd pathway in Aag2 cells. Importantly, the specificity of our RT-qPCR assays was verified by silencing genes involved in early signalling events within the Toll and Imd pathways as well as by silencing the downstream transcription factors. In addition, our luciferase reporter assays are based on transcription factor binding sites with previously verified specificity in *A. aegypti* [29]. Although we were able to develop REL1A-specific luciferase reporters, our REL2-responsive reporters were all also activated by REL1A. While the published NF-κB binding sites we based our reporters on specifically bound either REL1A or REL2 in an electrophoretic mobility shift assay (EMSA) [29], their specificity in *A. aegypti* cells was not reported. Other regulatory factors may therefore modulate the specificity of these NF-κB binding sites in Aag2 cells. Nevertheless, our REL2-responsive luciferase reporters still provide a specific read-out for the Imd pathway in Aag2 cells because we were unable to stimulate the Toll pathway under our experimental conditions.

All of the plasmids generated in this study have been deposited with Addgene (addgene.org).

### Non-responsiveness of the Toll pathway in Aag2 cells

In our hands, the Toll pathway was not induced by Gram-positive (*L. monocytogenes, S. aureus*) or Gram-negative (*E. coli*) bacteria, the fungal PAMP zymosan or the viral PAMP dsRNA in Aag2 cells. Our data expand on and are in agreement with Zhang *et al*., who made similar observations with *E. coli* using their RT-qPCR assays [42]. In addition, we and Zhang *et al*. found that even genes considered by some to be specific markers of the Toll pathway, such as defensins, appear to be regulated by the Imd pathway in Aag2 cells (Fig 1, 2) [42]. Furthermore, although Gram-positive bacteria and fungi are often said to be sensed by the Toll pathway while Gram-negative bacteria are said to be sensed by the Imd pathway, we found that all tested immune stimuli were capable of activating the Imd pathway in Aag2 cells. It is becoming more widely accepted that the regulation of AMP expression in *Diptera* is more complex than a model whereby individual genes are induced by specific stimuli through one defined NF-κB pathway under all conditions. For example, in *D. melanogaster*, certain AMPs have been shown to be primarily regulated through the Toll pathway in the fat body and through the Imd pathway in epithelia [18]. Cross-talk between NF-κB pathways has also been reported [19], and microarray analysis of genes induced after septic injury with *E. coli* or *Micrococcus luteus* revealed that most AMPs can be regulate by both the Toll and Imd pathways [15]. Our findings therefore further emphasise the need to carefully validate immune assays in each experimental system before drawing conclusions about specific pathways.

Why the Toll pathway appears to be non-responsive in Aag2 cells under our experimental conditions is unclear. We verified that all known signalling components of the Toll pathway downstream of SPÄTZLE are present in the published Aag2 reference genome (vectorbase.org) [50], and that there are no obvious mutations that would render pathway components non-functional. Of specific note, while the *PELLE* sequence in the Aag2 reference genome is suggestive of a premature stop codon, we used PCR and Sanger sequencing to verify that this potential premature stop codon is not present in the *PELLE* sequence of our Aag2 clone. We have noted such sequencing errors and misannotations in the *A. aegypti* and Aag2 reference genomes before [51,52]. While it is feasible that there is some real deficiency in the Toll pathway in Aag2 cells, we instead suspect that secreted proteins involved in the sensing of extracellular bacteria and activation of SPÄTZLE do not accumulate to sufficient levels in cell culture to facilitate Toll pathway activation. Similarly, the co-expression of SPÄTZLE alongside bacterial stimulation has been shown to be required for optimal Toll pathway activation in *D. melanogaster* S2 cells [43].

### Stimulation of REL2-regulated genes by viral PAMPs

Others have reported the sensing of virus-specific PAMPs by NF-κB pathways in *A. aegypti* [25]. Furthermore, DICER2 has been shown to sense infection with various RNA viruses in *D. melanogaster* and *Culex spp*. and *Aedes spp*. mosquitoes separately from its role in the RNAi pathway to stimulate the induction of *VAGO* via REL2 [46–49]. We extend these findings by demonstrating that a wide range of REL2-regulated genes are also induced by the general viral PAMP dsRNA and by CrPV infection in Aag2 cells. Our data indicate that dsRNA can be sensed by intracellular PRRs, however whether this sensing occurs via DICER2 in the context of REL2-regulated gene expression remains to be determined. Furthermore, the stimulation of REL2-regulated genes by treatment as well as by transfection with poly(I:C) is suggestive of additional extracellular or endosomal PRRs capable of detecting viral RNA in *A. aegypti*. This would mirror the situation in vertebrate systems where the endosomal membrane-bound TLR3 senses dsRNA [53]. A caveat to this interpretation of our data is that *D. melanogaster* cell cultures have been shown to take up dsRNA without transfection reagent [54], although this has not been reported for mosquito cell cultures.

In contrast to our data showing that CrPV infection induces AMP expression in Aag2 cells, arboviruses, including the flavivirus DENV and alphavirus Sindbis virus (SINV), do not induce AMP expression in Aag2 cells [40,41]. This is likely because flaviviruses and alphaviruses have evolved mechanisms to suppress immune signalling in their vector species [40,55]. In contrast, infection of Aag2 cells with CrPV is not a native virus-host combination and we hypothesise that CrPV therefore does not have the capacity to block REL2-regulated immune gene induction in Aag2 cells.

We have been careful here to distinguish between immune signalling through REL2 and the ‘classical’ Imd pathway characterised as a response to stimulation with bacterial PAMPs. Recently, gene expression by the *D. melanogaster* REL2 homologue RELISH has been shown to be stimulated by viral DNA through cGAS-STING signalling [56,57]. We therefore feel it is important to distinguish between the likely numerous and distinct signalling cascades converging on the transcription factor REL2 that respond to infection with different pathogens.

### Lack of *VAGO* induction by viral stimulation in Aag2 cells

We found no evidence for the induction of *VAGO1* or *VAGO2* in Aag2 cells following CrPV infection or stimulation with dsRNA. To our knowledge, *VAGO* induction in *A. aegypti* cell cultures has also not been reported by other groups. The *A. aegypti VAGO* promoter reportedly contains NF-κB binding sites [48], although it is unclear which *VAGO* gene this refers to. *VAGO1* has been shown to be induced *in vivo* in *A. aegypti* infected with DENV and is antiviral against DENV in Aag2 cells infected with *Wolbachia* [49]. Others observed the transient induction of *VAGO1*, but not *VAGO2*, in *A. aegypti* infected with yellow fever virus (YFV), but neither *VAGO1* nor *VAGO2* were induced by DENV or West Nile virus (WNV) [58]. In cell culture, the dicistrovirus *Drosophila* C virus (DCV) has been shown to induce *VAGO* in *D. melanogaster* S2 cells, while DENV induces *VAGO* in S2 and *Aedes albopictus* RML-12 cells and WNV induces *VAGO* in *Culex quinquefasciatus* Hsu cells [47–49]. It seems unlikely that Aag2 cells have a specific defect in factors required for *VAGO* induction given that we show here that other immune genes can be induced via REL2 by viral PAMPs. While VAGO expression may be stimulated in a virus-, cell type- or vector-specific manner, the overall picture from our and other’s data suggests that *VAGO* induction in *A. aegypti* is not a robust or consistent general response to viral infection.

Overall, our highly specific and well characterised tools for studying the Imd pathway in Aag2 cells, and our observation that dsRNA can induce signalling through REL2, pave the way for future studies defining molecular virus-immune interactions in *A. aegypti* that could ultimately inform the development of transmission-incompetent vectors to reduce the global arboviral disease burden.

